# Resistance to pentamidine is mediated by AdeAB, regulated by AdeRS, and influenced by growth conditions in *Acinetobacter baumannii* ATCC 17978

**DOI:** 10.1101/265520

**Authors:** Felise G. Adams, Uwe H. Stroeher, Karl A. Hassan, Shashikanth Marri, Melissa H. Brown

## Abstract

In recent years, effective treatment of infections caused by *Acinetobacter baumannii* has become challenging due to the ability of the bacterium to acquire or up-regulate antimicrobial resistance determinants. Two component signal transduction systems are known to regulate expression of virulence factors including multidrug efflux pumps. Here, we investigated the role of the AdeRS two component signal transduction system in regulating the AdeAB efflux system, determined whether AdeA and/or AdeB can individually confer antimicrobial resistance, and explored the interplay between pentamidine resistance and growth conditions in *A. baumannii* ATCC 17978. Results identified that deletion of *adeRS* affected resistance towards chlorhexidine and 4’,6-diamidino-2-phenylindole dihydrochloride, two previously defined AdeABC substrates, and also identified an 8-fold decrease in resistance to pentamidine. Examination of Δ*adeA*, Δ*adeB* and Δ*adeAB* cells augmented results seen for Δ*adeRS* and identified a set of dicationic AdeAB substrates. RNA-sequencing of Δ*adeRS* revealed transcription of 290 genes were ≥2-fold altered compared to the wildtype. Pentamidine shock significantly increased *adeA* expression in the wildtype, but decreased it in Δ*adeRS*, implying that AdeRS activates *adeAB* transcription in ATCC 17978. Investigation under multiple growth conditions, including the use of Biolog phenotypic microarrays, revealed resistance to pentamidine in ATCC 17978 and mutants could be altered by bioavailability of iron or utilization of different carbon sources. In conclusion, the results of this study provide evidence that AdeAB in ATCC 17978 can confer intrinsic resistance to a subset of dicationic compounds and in particular, resistance to pentamidine can be significantly altered depending on the growth conditions.

## Introduction

*Acinetobacter baumannii* causes a range of disease states including hospital-acquired pneumonia, blood stream, urinary, wound and bone infections, and is responsible for epidemic outbreaks of infection worldwide [1]. Such infections are often very difficult to treat due to the multidrug resistant (MDR) character of isolates displayed by this organism [2, 3]. In addition to the impressive propensity of the organism to acquire genetic elements carrying resistance determinants [2, 4, 5], up-regulation resulting in overproduction of resistance nodulation cell-division (RND) drug efflux systems through integration of insertion sequence elements or mutations in regulatory genes, has also been deemed a major contributor to the MDR phenotype [6-9]. The best studied RND efflux systems in *A. baumannii* include AdeABC [10], AdeFGH [11] and AdeIJK [12]. Of particular interest is the AdeABC system which affords resistance to diverse antibiotics, biocides and dyes [10, 13-15], and has gained attention due to its high incidence of over-expression across many MDR *A. baumannii* clinical isolates, primarily from incorporation of point mutations in the genes encoding its positive regulator, AdeRS [6, 8, 13, 16]. Typically RND pumps consist of three proteins that form a complex; the absence of any of these components renders the entire complex non-functional [17]. Interestingly, deletion of *adeC* in the *A. baumannii* strain BM4454 did not affect resistance towards two substrates of the pump suggesting that AdeAB can utilize an alternative outer membrane protein (OMP) to efflux antimicrobial compounds [13].

The genetic arrangement of the AdeABC system places *adeABC* in an operon that is divergently transcribed to the regulatory *adeRS* two component signal transduction system (TCSTS). Expression of *adeABC* occurs by binding of AdeR to a ten base-pair direct repeat motif found within the intercistronic region separating these operons [18, 19]. Many clinical *A. baumannii* isolates harbor different genetic arrangements of the *adeRS* and *adeABC* operons [20], and whether regulation via AdeRS is conserved in these strains is not completely understood.

With an increase in infections caused by MDR isolates across many bacterial species, including *A. baumannii,* understanding mechanisms of resistance and how resistance to and evasion of treatments has evolved over time has become a key research topic. Furthermore, determining the impact of expression of resistance determinants within the host environment and its effect on the efficacy of therapeutic treatments has gained attention. For example, when *Pseudomonas aeruginosa* is grown using L-glutamate as the sole carbon source, resistance to the related compounds polymyxin B and colistin increased ≥ 25-and 9-fold, respectively [21]. Other studies have shown that the bioavailability of cations such as iron can have a drastic effect towards the resistance of a number of antimicrobials across a range of pathogenic bacterial species [22]. Despite *A. baumannii* being recognized as a major human pathogen, these types of studies are limited for this organism.

This study aimed to determine the regulatory role of the AdeRS TCSTS in *A. baumannii* ATCC 17978, a clinical isolate which only encodes the *adeAB* subunits and identify whether AdeA and or AdeB alone can confer antimicrobial resistance in the ATCC 17978 background. Phenotypic characterization of a constructed panel of deletion strains identified that alterations to the *adeRS* and *adeAB* operons of ATCC 17978 reduced resistance to a subset of dicationic compounds, including pentamidine. As a recent study highlighted the effectiveness of pentamidine in combination therapy to treat infections caused by Gram-negative pathogens [23], we sought to further examine alternative mechanisms of pentamidine resistance in *A. baumannii.* The type of carbon source and availability of iron were identified as affecting pentamidine resistance, thereby revealing interconnectedness between metabolic and resistance strategies within this formidable pathogen.

## Materials and methods

### Bacterial strains, plasmids and growth conditions

*A. baumannii* ATCC 17978 [24] was obtained from the ATCC and is designated as wildtype (WT). Bacterial strains, plasmids and primers used in this study are summarized in S1 and S2 Tables, respectively. Bacterial strains were cultured using Luria-Bertani (LB) broth or LB agar plates, under aerobic conditions at 37°C, unless otherwise stated. Antibiotic concentrations used for selection were; ampicillin 100 mg/L, erythromycin (ERY) 25 mg/L, tetracycline 12 mg/L and gentamicin (GEN) 12.5 mg/L. All antimicrobial agents were purchased from Sigma with the exception of ampicillin which was purchased from AMRESCO.

M9 minimal medium agar plates were generated using a stock solution of 5 × M9 salts (200 mM Na_2_HPO_4_.7H_2_O, 110 mM KH_2_PO_4_, 43 mM NaCl and 93 mM NH_4_O) and subsequently diluted 1:5 and supplemented with 2 mM MgSO_4_, 0.1 mM CaCl_2_ and 0.4% (w/v) of various carbon sources on the day of use. M9 minimal medium was supplemented with 0.4% (w/v) of glucose, or succinic, fumaric, citric, α-ketoglutaric, pyruvic, glutamic or oxaloacetic acids. All carbon compounds were purchased from Sigma with the exception of citric and pyruvic acids which were purchased from Chem-Supply (Gillman, Australia). For solid media, J3 grade agar (Gelita, Australia) was used at a final concentration of 1%.

Iron-chelated and iron-rich conditions were achieved by the addition of dipyridyl (DIP) or FeSO4.7H_2_O (Scharlau) at final concentrations of 100 and 200 µM or 2.5 and 5 mM in Mueller-Hinton (MH) agar, respectively.

### Antimicrobial susceptibility testing

The resistance profiles of WT and derivative strains were determined via broth microdilution methods [25] which were performed in duplicate with a minimum of three biological replicates using bacteria cultured in MH broth. Growth was monitored at 600 nm (OD_600_) using a FLUOstar Omega spectrometer (BMG Labtech, Germany) after overnight incubation at 37°C.

For disk diffusion assays, strains were grown to mid-log phase in MH broth, diluted to OD600 = 0.1 in fresh MH broth and 100 µL plated onto MH agar plates on which antimicrobial loaded filter disks were overlaid. Alternatively, when assessing zones of clearing using M9 minimal medium agar, strains were washed three times in phosphate buffered saline (PBS) and diluted to OD600 = 0.1 in fresh PBS. Tested antimicrobials were pentamidine isethionate, chlorhexidine dihydrochloride and 4’,6-diamidino-2-phenylindole dihydrochloride (DAPI) at the final concentrations of 125, 2.5 and 1.6 μg, respectively. After overnight incubation, callipers were used to measure the diffusion distances which were determined as half of the inhibition zone diameter, less the length of the disk. At least three independent experiments in duplicate were undertaken. Statistical significance was determined using the Student’s t-test (two-tailed, unpaired) and *P* values <0.0005 were considered significant.

For plate dilution assays, strains were grown to mid-log phase in MH broth, washed three times in PBS and diluted to OD_600_ = 0.1 in fresh PBS. Serial dilutions (10-fold) were prepared of which 5 μl was spotted onto M9 minimal or MH agar containing various concentrations of pentamidine (32, 64, 128, 256 and 512 mg/L) and growth assessed after overnight incubation at 37°C. To test the effect of growth of different carbon sources, M9 minimal media was supplemented with 0.4% (w/v) of succinic, fumaric, citric, α-ketoglutaric or oxaloacetic acids.

### Construction of *A. baumannii* ATCC 17978 deletion strains and genetic complementation

Two methods were adopted to generate specific gene deletions in *A. baumannii* ATCC 17978. The first method used to generate Δ*adeRS* utilized a sacB-based strategy [26], where genetic sequence flanking the region of interest and the ERY resistance cassette from pVA891 [27] were PCR amplified, purified and subsequently cloned into a modified pBluescript SK+ II vector that had the *Bam*HI site removed by end-filling, generating pBl_adeRS. Confirmed clones were re-cloned into *Xba*l-digested pEX18Tc [28]. The resulting pEX18Tc_adeRS vector was introduced into ATCC 17978 cells via electroporation, as previously described [29]. Transformants were selected on LB agar supplemented with ERY. Counter-selection was undertaken on M9 minimal agar containing ERY and 5% sucrose. The second method used to generate *ΔadeA, ΔadeB* and *ΔadeAB* strains utilized the RecET recombinase system with modifications [30]. Briefly, 400-600 bp of sequence flanking the gene(s) were used as templates in a nested overlap extension PCR with the ERY resistance cassette from pVA891 [27]. Approximately, 3.5-5 μg of the purified linear PCR product was electroporated into ATCC 17978 cells harboring the vector pAT04 and recovered as previously described [30]. Recombinants were selected on LB agar supplemented with ERY. All mutants generated in this study were confirmed by PCR amplification and Sanger sequencing. Primers utilized to generate mutant strains are listed in S2 Table.

For genetic complementation of mutant strains, WT copies of *adeRS* and *adeAB* were cloned into pWH1266 [31] where transcription was driven by the tetracycline promoter. Resulting plasmids denoted pWH::adeRS and pWH::adeAB, respectively were confirmed by Sanger sequencing. The GEN resistance cassette from pUCGM [32] was PCR amplified and cloned into *BamHI* digested pWH: *:adeAB* generating pWHgent::adeAB thus abrogating transcription from the pWH1266 tetracycline promoter. Plasmids were introduced into appropriate *A. baumannii* cells as previously described [33]. Primers used to generate complementation vectors are listed in S2 Table.

### Cell treatments and RNA isolation

RNA was isolated from WT and *ΔadeRS* cells and Hiseq RNA transcriptome analysis performed following methodologies as outlined previously [34].

For pentamidine stress assays, WT and *ΔadeRS* strains were grown overnight in MH broth, sub-cultured 1:100 in fresh medium and grown to OD_600_ = 0.6; they were subsequently split into two 10 mL cultures. One 10 mL sample was treated with 7.8 mg/L of pentamidine (0.5 x MIC for *ΔadeRS),* whilst the other remained untreated. Cultures were grown for an additional 30 min before total RNA was extracted as outlined previously [34].

### Bioinformatic analysis

Bioinformatic analysis of RNA-seq data was undertaken as described previously [34], with the modification that obtained reads were mapped to the recently re-sequenced *A. baumannii* ATCC 17978 genome (GenBank: CP012004). RNA-seq data have been deposited in the gene expression omnibus database, accession number GSE102711 (https://www.ncbi.nlm.nih.gov/geo/query/acc.cgi?acc=GSE102711).

### Quantitative Real-Time PCR

Pentamidine stress and RNA-seq validation experiments were achieved using a two-step qRT-PCR method. RNA samples were first purified as previously described [34], subsequently DNaseI treated (Promega) and then converted to cDNA using an iScript™ cDNA synthesis kit (Biorad), following the manufacturer’s instructions. The cDNA generated was used as a template for qRT-PCR using the SYBR^®^ Green JumpStart™ Taq readymix™ (Sigma) in a 20 μl final volume. Either a Rotor-Gene Q (Qiagen, Australia) or RG-3000 (Corbett Life Science, Australia) instrument was used for quantification of cDNA using the following protocol; 1 min at 95°C, followed by 40 cycles of 10 sec at 95°C, 15 sec at 57°C and 20 sec at 72°C. Melt curve analyses were undertaken to ensure only the desired amplicon was generated. Primers used (S2 Table) for amplification of cDNA transcripts were designed using NetPrimer (www.premiersoft.com). Transcriptional levels for RNA-seq validation experiments were corrected to *GAPDH* levels prior to normalization to the ATCC 17978 WT. For pentamidine stress experiments, transcriptional levels of *adeA* were corrected to 16S rDNA levels prior to being normalized to their respective untreated *A. baumannii* cultures. Transcriptional variations were calculated using the 2^−ΔΔCT^ method [35]. Results for pentamidine stress experiments display the mean Log_2_ fold change (± SEM) of three biological replicates each undertaken in triplicate. Statistical analyses were performed by Student’s t-test, twotailed, unpaired; ** = *P* <0.01 and *** = *P* <0.001.

### Phenotypic microarray analysis

*A. baumannii ΔadeRS* cells were cultured on LB agar overnight at 37°C. A suspension of cells was made from a single colony in Biolog IF-0 inoculation fluid (Biolog, Inc.) to 85% transmittance and was subsequently diluted 1:200 in Biolog IF-0 containing dye A (Biolog, Inc.) and 0, 8, 16, 32 or 64 mg/L pentamidine. One hundred µL of each dilution was added to each well of the Biolog PM01 and PM02A MicroPlates and placed in an Omnilog automatic plate reader (Biolog, Inc.) for 72 h at 37°C. Color formation from the redox active dye (Biolog dye A) was monitored every 15 min. Data obtained from respiration of *ΔadeRS* under different pentamidine conditions were individually overlaid against the untreated control using the OmniLog File Management/Kinetic Analysis software v1.20.02, and analyzed using OmniLog Parametric Analysis software v1.20.02 (Biolog, Inc.).

## Results

### Generation of *ΔadeRS* and complemented strains

The *adeRS* operon of *A. baumannii* ATCC 17978 was disrupted through the introduction of an ERY resistance cassette to produce a *ΔadeRS* strain. Complementation of *ΔadeRS* was achieved via reintroduction of a WT copy of *adeRS in trans* on the pWH1266 shuttle vector, generating *ΔadeRS pWH: :adeRS* and comparisons were made to cells carrying the empty vector control (*ΔadeRS* pWH1266). To ensure deletion of *adeRS* did not affect viability, growth was monitored by OD_600_ readings in MH broth over an 8 h period; results identified no significant perturbations in the growth rate in laboratory media compared to WT cells (data not shown).

### Transcriptomic profiling of *ΔadeRS*

Transcriptome profiling by RNA-sequencing (RNA-seq) of RNA isolated from *ΔadeRS* compared to that from WT ATCC 17978 identified 290 differentially expressed (≥ 2-fold) genes; 210 up-regulated and 80 down-regulated (Fig 1 and S3 Table). RNA-seq results were confirmed by quantitative Real-Time PCR (qRT-PCR) for nine genes that displayed different levels of expression in *ΔadeRS* compared to WT; a good correlation between methods was observed (S1 Fig).

**Fig 1.**
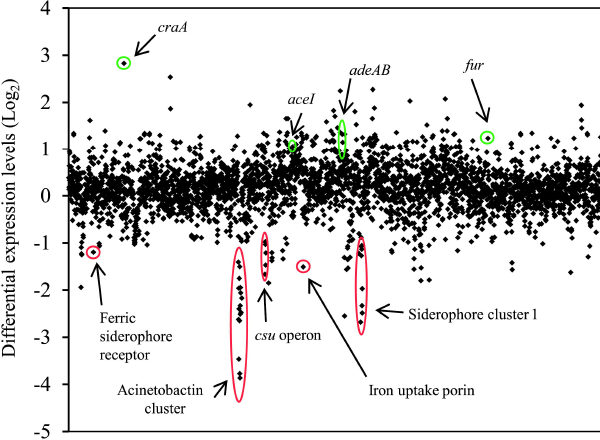
Global transcriptomic response differences of A. baumannii ATCC 17978 after deletion of adeRS. Each diamond marker represents a predicted gene within the genome ordered according to locus tag along the X-axis and differential expression of WT against *ΔadeRS* are displayed on the Y-axis (Log_2_). Positive and negative Log_2_-values correlate to up- and down-regulated genes, respectively. Green and red circles highlight genes/gene clusters of interest that have been up- and down-regulated, respectively. See S3 Table for the full list of genes that were differentially expressed ≥1 Log_2_ fold.

Expression of a number of efflux proteins was affected by inactivation of *adeRS.* For example, the *craA* major facilitator superfamily transporter (ACX60_01760) shown to confer resistance to chloramphenicol and recently chlorhexidine efflux [36] was the highest up-regulated gene (7.1-fold; Fig 1) and *adeA* (ACX60_09125), *adeB* (ACX60_09130) and *aceI* (ACX60_07275) which encode the AdeAB and the AceI efflux pumps, were up-regulated by 2.5, 1.43, and 2-fold, respectively, following deletion of *adeRS.*

Expression of multiple genes involved in virulence were down-regulated in the *ΔadeRS* strain, including the type 1 pilus operon *csuA/BABCDE* (ACX60_06480-06505) (3.2- to 2-fold) and the siderophore-mediated iron-acquisition systems acinetobactin (ACX60_05590-05680) and siderophore 1 (ACX60_09665-09720) clusters (7.7- to 1.1-fold and 5.4- to 0.3-fold, respectively). In contrast, the ferric uptake regulator gene *(fur*) (ACX60_13910) was up-regulated 2.4-fold. Despite these alterations in expression levels of iron siderophore clusters and their regulator, no significant growth defects were identified when *ΔadeRS* was grown in the presence of 200 μM of DIP (data not shown).

### Deletion of *adeRS* in ATCC 17978 reduced susceptibility to a limited number of antimicrobial agents

To assess if changes in expression of the *adeAB* and *craA* drug efflux genes identified in the transcriptome of the *ΔadeRS* derivative translated to an alteration in resistance profile, antibiogram analyses were undertaken. Resistance to a number of antibiotics including those that are known substrates of the CraA and AdeABC pumps were assessed. Surprisingly, despite *craA* being the highest up-regulated gene, no change in resistance to the primary substrate of CraA, chloramphenicol [36], was seen (data not shown). Previous studies examining the level of antimicrobial resistance conferred by AdeABC indicate that only when deletions are generated in strains which overexpress AdeABC is there a significant impact on the antibiogram [10, 13, 15, 16, 37], implying that AdeABC confers only minimal to no intrinsic resistance. This was supported in our analysis as the MIC for tetracycline, tigecycline, GEN, kanamycin, nalidixic acid, ampicillin, streptomycin and amikacin, all previously identified AdeABC substrates, remained unchanged, whilst resistance to norfloxacin and ciprofloxacin increased 2-fold (data not shown) and chlorhexidine decreased 2-fold for the *ΔadeRS* mutant compared to WT (Table 1).

**Table 1.** Antibiotic susceptibility of *A. baumannii* ATCC 17978, deletion mutants and complemented strains.

| Strain | MIC (mg/L)<sup>ab</sup> |  |  |
| --- | --- | --- | --- |
|  | PENT | DAPI | CHX |
| WT | 125 | 4 | 8 |
| $\Delta adeRS$ | **15.6** | **1** | **4** |
| $\Delta adeAB$ | **15.6** | **1** | **4** |
| $\Delta adeA$ | **15.6** | **1** | **4** |
| $\Delta adeB$ | **15.6** | **1** | **4** |
| $\Delta adeRS$ pWH::adeRS | 125 | 1 | 8 |
| $\Delta adeRS$ pWH1266 | 15.6 | 0.5 | 4/2 |
| $\Delta adeRS$ pWH::adeAB | 31.3 | 4 | 4 |
| $\Delta adeRS$ pWHgent::adeAB | 15.6 | 1 | 4 |
| $\Delta adeRS$ pWHgent | 7.8 | 0.5 | 4 |
| *ΔadeA* pWHgent::*adeAB* | 62.5 | 2 | 8/4 |
| *ΔadeA* pWHgent | 7.8 | 0.5 | 4 |
| *ΔadeB* pWHgent::*adeAB* | 62.5 | 4 | 4 |
| *ΔadeB* pWHgent | 7.8 | 1 | 2 |
| *ΔadeAB* pWHgent::*adeAB* | 62.5 | 4 | 4 |
| *ΔadeAB* pWHgent | 7.8 | 1 | 2 |

<sup>a</sup>PENT, pentamidine; DAPI, 4',6-diamidino-2-phenylindole; CHX, chlorhexidine.
<sup>b</sup>Values highlighted in bold indicate MIC values altered two-fold or greater compared to WT ATCC 17978.

Other compounds tested included; colistin, polymyxin B, rifampicin, triclosan, novobiocin, benzalkonium, methyl viologen, pentamidine, DAPI and dequalinium. Of these, significant differences in the MICs of *ΔadeRS* versus WT were observed only for the diamidine compounds, pentamidine and DAPI, where an 8- and 4-fold reduction in resistance was identified, respectively (Table 1). Thus, taken together the *ΔadeRS* strain showed reduced resistance to the bisbiguanide chlorhexidine, and the diamidines pentamidine and DAPI which display structural similarities [38] (S2 Fig). Resistance to these substrates was partially or fully restored by complementation using *ΔadeRS pWH::adeRS* (Table 1).

### Both AdeA and AdeB are required for intrinsic antimicrobial resistance in ATCC 17978

Since the 10-bp direct repeat where AdeR has been demonstrated to bind in other *A. baumannii* strains [18, 19] is also present in the ATCC 17978 *adeA-adeR* intercistronic region (data not shown), we hypothesized that the decreased resistance towards the subgroup of dicationic compounds seen in *ΔadeRS* resulted from changes in *adeAB* expression. To test this, deletion strains targeting *adeAB,* as well as individual *adeA* and *adeB* mutations were generated in ATCC 17978.

Using *ΔadeAB,* MIC analyses verified that deletion of the pump resulted in negligible changes in resistance to a subset of the antimicrobials tested for *ΔadeRS* (data not shown). However, an identical resistance pattern for the dicationic compounds as that afforded by *ΔadeRS* was observed (Table 1). Complementation of the inactivated genes partially restored resistance to all compounds, validating that AdeAB plays a direct role in resistance to these dicationic compounds (Table 1).

### AdeRS is critical for increased expression of *adeAB* following pentamidine exposure

From MIC analysis of the ATCC 17978 derivatives, it was proposed that the presence of the AdeRS TCSTS increased *adeAB* expression consequently providing resistance to pentamidine, chlorhexidine and DAPI (Table 1). To confirm this, the level of *adeA* transcription was assessed by qRT-PCR of RNA isolated from WT and *ΔadeRS* strains after addition of a sub-inhibitory concentration of pentamidine. Transcription of the *adeAB* operon was significantly up-regulated in WT and down-regulated in *ΔadeRS* following pentamidine stress (Fig 2). Additionally, qRT-PCR was used to determine if *adeS* expression levels altered after pentamidine stress in WT cells. It was found that *adeS* expression increased less than <2-fold compared to untreated WT cells (data not shown). To phenotypically support the transcriptional evidence that *adeRS* initiates transcription of the ATCC 17978 *adeAB* operon, additional antibiograms were determined. Using the shuttle vector pWH1266, two clones were constructed; *pWH/.adeAB* and pWHgent:*:adeAB,* where the GEN resistance cartridge cloned in the latter vector inhibited transcription of *adeAB* from the tetracycline promoter naturally present in pWH1266. MIC analyses determined that the carriage of pWHgent::adeAB in *ΔadeRS* did not differ from results obtained for *ΔadeRS.* Conversely, *ΔadeRS* cells with pWH::adeAB displayed a 4-and 8-fold increase in pentamidine and DAPI resistance, respectively (Table 1). Collectively, these results suggest that expression of *adeAB* and subsequent resistance to the dicationic compounds in ATCC 17978 can only occur when AdeRS is present.

**Fig 2.**
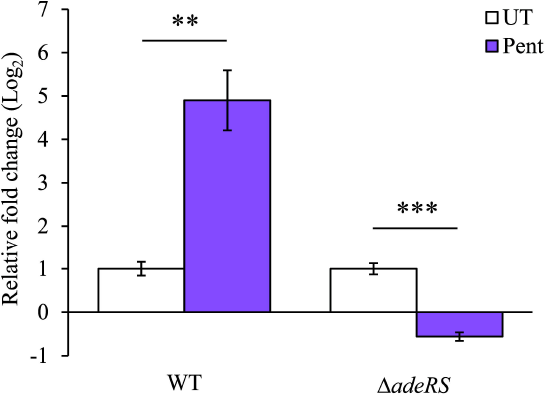
Increased expression of adeA following pentamidine stress is dependent on the presence of AdeRS in ATCC 17978. Transcriptional levels of *adeA* (ACX60_09125) from WT and *ΔadeRS* were determined by qRT-PCR after 30 min shock with 7.8 mg/L of pentamidine (Pent) (0.5 x MIC of *AadeRS)* and corrected to untreated cells (UT) after normalization to 16S. Bars represent the mean fold change (Log2) of three biological replicates undertaken in triplicate, and error bars represent ± SEM. Statistical analyses were performed by Student’s t-test, two-tailed, unpaired; ** = *P* < 0.01 and *** = *P* < 0.001.

### Carbon source utilization alters resistance to pentamidine

Pentamidine, a drug known to be effective in the treatment of fungal and protozoan infections has gained recent attention in a bacterial context. Pentamidine has shown synergy with Gram-positive antibiotics, potentiating their activity towards Gram-negative bacteria [23]. As such, pentamidine has been proposed to be utilized as an adjunct therapy for MDR bacteria, including *A. baumannii* [23]. To identify additional pentamidine resistance mechanisms employed by *A. baumannii,* growth in different media was assessed. Disk diffusion assays identified that in *A. baumannii,* resistance to pentamidine was affected by the carbon source provided in M9 minimal medium (S4 Table), whilst for chlorhexidine and DAPI this pattern was not conserved indicating a pentamidine-specific response (data not shown). To determine the MIC levels for pentamidine, plate dilution experiments were undertaken for WT, *ΔadeRS* and *ΔadeAB* strains provided with varied carbon sources (Fig 3). Growth of *ΔadeRS* and *ΔadeAB* cells on MH agar were significantly perturbed at 32 mg/L of pentamidine. This MIC drastically differs when succinic acid was utilized as the sole carbon source, as growth was maintained up to 512 mg/L of pentamidine for all strains tested (Fig 3). Fumaric, α-ketoglutaric and oxaloacetic acids also increased pentamidine resistance by 8-fold for *ΔadeRS* and *ΔadeAB* strains when compared to the MIC obtained for MH agar. Growth of *ΔadeRS* and *ΔadeAB* mutants was inhibited at a higher dilution factor in the presence of oxaloacetic acid compared to fumaric and α-ketoglutaric acids whilst growth using citrate as the sole carbon source negatively affected resistance to WT cells, decreasing resistance 2-fold.

**Fig 3.**
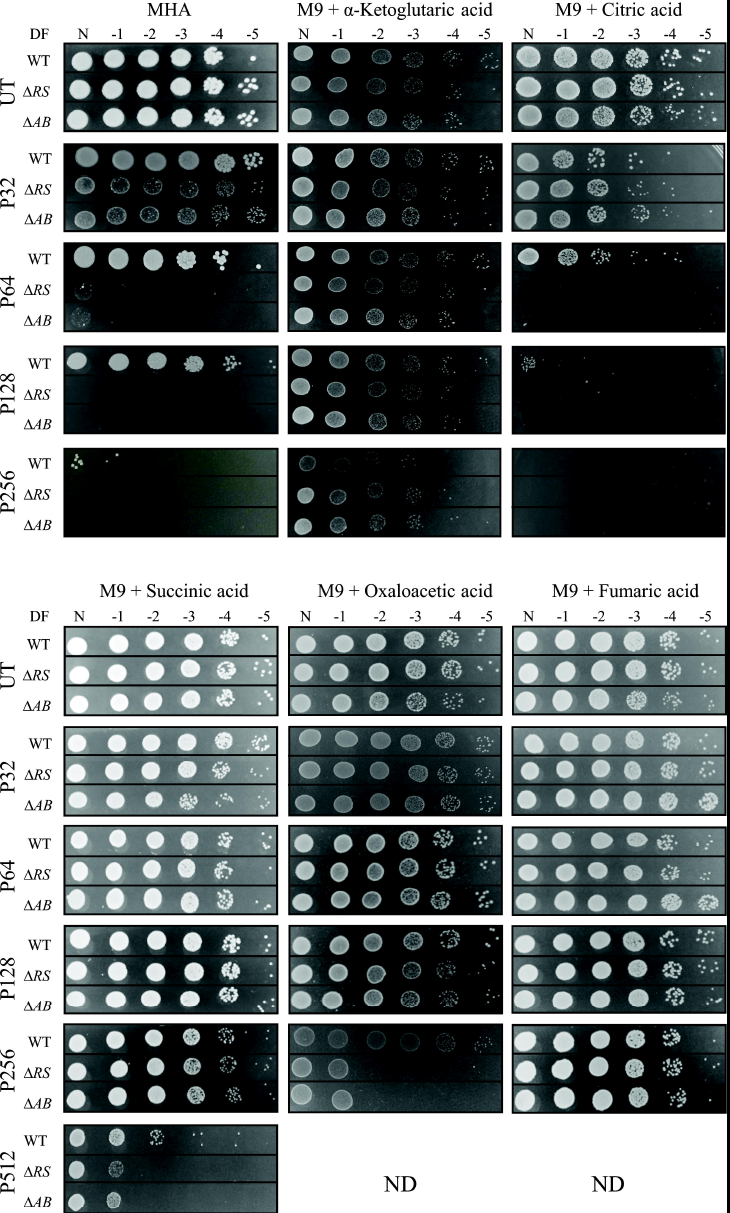
Resistance to pentamidine is modulated by carbon sources available in the growth medium. Ten-fold serial dilutions of *A. baumannii* ATCC 17978 (WT), *ΔadeRS (ΔRS)* and *ΔadeAB (ΔAB)* cells were used to compare the concentration of pentamidine that inhibits growth in different media, Mueller-Hinton agar (MHA) was used as a comparative control. Images display serial 1:10 dilutions after overnight incubation at 37°C, where DF is abbreviated for dilution factor and N represents undiluted cells. Strains were grown in the absence of pentamidine (UT) or presence of 32, 64, 128, 256 and 512 mg/L of pentamidine (P32, P64, P128, P256, and P512, respectively). Carbon sources tested in M9 minimal medium were used at a final concentration of 0.4% (w/v). ND, not done due to precipitation of pentamidine once added into the molten medium. Figures are representative examples of results obtained.

Biolog phenotypic arrays were undertaken to identify additional synergistic or antagonistic relationships between carbon sources and pentamidine resistance. Respiration of *ΔadeRS* cells for a total of 190 carbon compounds was assessed at various pentamidine concentrations (S3 Fig). From this, an additional ten compounds were identified that increased resistance to pentamidine in *ΔadeRS* at 64 mg/L (Fig 4). Despite minimal changes in pentamidine resistance in the presence of citric acid for *ΔadeRS* (Fig 3), this compound negatively affected respiration at lower pentamidine concentrations (S3 Fig). Surprisingly, succinic, α-ketoglutaric and fumaric acids failed to restore respiration at 64 mg/L of pentamidine for *ΔadeRS* (S3 Fig). By extending the incubation period for another 72 h, respiration in the presence of succinic or fumaric acids was restored to levels similar to the untreated control (data not shown). This may indicate that succinic and fumaric acids significantly lag in their ability to recover the cells from pentamidine in the IF-0 medium (Biolog Inc.) whilst α-ketoglutaric acid recovery is dependent on growth medium.

**Fig 4.**
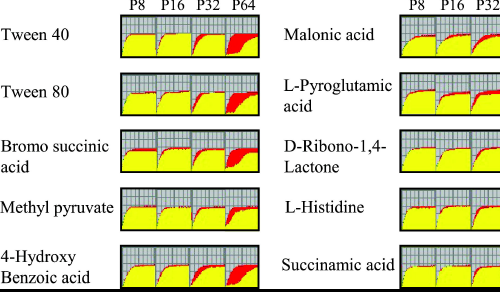
Kinetic response curves paralleling bacterial growth from Biolog PM01 and PM2A plates identify ten carbon sources that increase pentamidine resistance in *ΔadeRS*. P8, P16, P32 and P64 represent kinetic response curves at 8, 16, 32, or 64 mg/L of pentamidine compared to the untreated control, respectively. Red curves represent respiration of untreated *ΔadeRS,* whilst respiratory activity which overlaps between the control and the sample in the different experimental conditions is represented in yellow. Only carbon compounds that promote at least 50% maximal respiration and induce a recovery response by 36 h are shown. See S3 Fig for respiration curves for all tested treatments.

### Bioavailability of iron correlates with pentamidine resistance

Iron has been shown to influence pentamidine resistance in protozoan species [39], thus we assessed if its plays a similar role in *A. baumannii.* The addition of ferrous sulphate to the growth medium significantly reduced the zone of clearing from pentamidine in a dose-dependent manner up to a final concentration of 5 mM for WT and mutant strains (Table 2 and S4 Fig). Furthermore, chelation of iron using DIP resulted in WT cells becoming more susceptible to pentamidine compared to the untreated control (Table 2). This modified pentamidine susceptibility is iron specific as inclusion of other cations in the growth medium (zinc, copper, manganese, cobalt, nickel) did not significantly affect the zones of clearing (data not shown). Using inductively-coupled plasma mass spectrometry the internal iron concentration in WT and *ΔadeRS* cells was determined in the absence/presence of a sub-inhibitory concentration of pentamidine. Internal iron concentrations remained essentially unchanged in both strains under the different conditions tested (data not shown), indicating that iron/pentamidine interactions may occur outside of the cell and the observed response was not due to a reduced capacity to store iron.

**Table 2.** Zones of clearing for *A. baumannii* ATCC 17978 and deletion mutants grown on Mueller-Hinton agar with the addition or chelation of iron after pentamidine exposure.

| Strain | Zone of clearing (mm)<sup>ab</sup> |  |  |  |  |
| --- | --- | --- | --- | --- | --- |
| | WT | $\Delta adeRS$ | $\Delta adeAB$ | $\Delta adeA$ | $\Delta adeB$ |
| **Medium<sup>c</sup>** |  |  |  |  |  |
| MHA control | 1.4 ± 0.3 | 6.9 ± 0.5 | 6.9 ± 0.3 | 6.9 ± 0.4 | 7.2 ± 0.4 |
| + 2.5 mM FeSO<sub>4</sub> | **0.8 ± 0.1<sup>\*</sup>** | **5.4 ± 0.2<sup>\*</sup>** | **5.6 ± 0.3<sup>\*</sup>** | **5.4 ± 0.1<sup>\*\*\*</sup>** | **5.6 ± 0.1<sup>\*\*\*</sup>** |
| + 5 mM FeSO<sub>4</sub> | **0.7 ± 0.2<sup>\*</sup>** | **2.5 ± 0.8<sup>\*\*\*</sup>** | **2.8 ± 0.3<sup>\*\*\*</sup>** | **3.2 ± 0.3<sup>\*\*\*</sup>** | **2.9 ± 0.6<sup>\*\*\*</sup>** |
| + 7.5 mM FeSO<sub>4</sub> | NG | NG | NG | NG | NG |
| + 100 µM DIP | 1.7 ± 0.1 | 6.7 ± 0.2 | 6.9 ± 0.1 | 6.2 ± 0.2 | 6.6 ± 0.2 |
| + 200 µM DIP | **5 ± 0.3<sup>\*\*\*</sup>** | 6.7 ± 0.4 | 6.6 ± 0.4 | 6.1 ± 0.1 | 6.4 ± 0.1 |
| **pentamidine exposure.** |  |  |  |  |  |

<sup>a</sup>Averaged values are displayed ± SD. Statistical analyses were performed by Student's *t*-test, two-tailed, unpaired; \* = *P* < 0.0005 and \*\*\* = *P* < 0.000001.
<sup>b</sup>Values given in bold indicate a significant difference between a strain grown under iron-rich or iron-limited conditions versus its respective MHA control.
<sup>c</sup>MHA, Mueller-Hinton agar; FeSO<sub>4</sub>, ferrous sulphate; DIP, 2',2' dipyridyl; NG, no growth due to iron toxicity.

## Discussion

The *adeRS* and *adeABC* operons of *A. baumannii* have gained considerable attention due to their role in regulating and conferring multidrug resistance, respectively [8, 13, 16, 40]. Multiple genetic arrangements of the *adeABC* operon in *A. baumannii* clinical isolates exist, where 35% of 116 diverse isolates lack the *adeC* OMP component [20]. This is also the case for the clinical isolate *A. baumannii* ATCC 17978, which was chosen for further analysis in this study. It is not uncommon for RND pumps to recruit an alternative OMP to form a functional complex [41-43]. This may also be the situation for AdeAB in *A. baumannii* ATCC 17978, since in an *Escherichia coli* background, *adeAB* can be co-expressed with *adeK,* an OMP belonging to the *A. baumannii* AdeIJK RND complex to confer resistance [44]. However, whether AdeB can work alone or function with alternative membrane fusion proteins such as *adeI* or *adeF* is not known. As the resistance profile of *ΔadeA* was indistinguishable from the *ΔadeAB* and *ΔadeB* strains, it provides confirmatory evidence that both *adeA* and *adeB* are required for efficient efflux to only a subset of dicationic compounds in ATCC 17978. Further assessments will be required to verify if this phenotype is maintained across other *A. baumannii* isolates and extends to other structurally-similar compounds.

It has been found previously, that the introduction of shuttle vectors expressing WT copies of genes *in trans* do not restore resistance profiles back to WT levels as also seen in this study [45-48]. Using qRT-PCR, *adeS* was expressed at low levels in WT cells even after pentamidine shock (data not shown). Complementation studies with *adeRS* thus could be influenced by copy number effects, which may perturb the native expression levels of these proteins within the cell. Additionally, the differing modes of action of the dicationic compounds may have influenced the MIC results, as fold-shifts in resistance were observed even when the empty pWH1266 vector was being expressed (Table 1).

The transcriptomic analysis of ATCC 17978 Δ*adeRS* revealed changes in gene expression were not limited to *adeAB* but included many genes, some of which have been shown to be important in *A. baumannii* pathogenesis [49, 50] and virulence *in vivo* [51]. Although this had been seen before in an *adeRS* mutant generated in the MDR *A. baumannii* isolate AYE [16], here the transcriptional changes were to a largely different subset of genes. In particular, deletion of *adeRS* in AYE resulted in decreased expression of *adeABC* by 128-, 91-, and 28-fold, respectively [16]. This decreased gene expression was expected as AYE naturally contains a point mutation in AdeS (producing an Ala 94 to Val substitution in AdeS) resulting in constitutive expression of *adeABC,* thus contributing to its MDR phenotype [2, 16]. Conversely, our transcriptome data indicated that *adeAB* is only expressed at low levels in WT ATCC 17978, a strain labelled as having a ‘drug susceptible’ phenotype [24].

Currently, the environmental signal(s) that interact with the periplasmic sensing domain of AdeS are not known. Antimicrobial compounds can directly stimulate autophosphorylation of specific TCSTS which in turn directly regulate the expression of genes providing resistance to that compound [52, 53]. It is unlikely that pentamidine is the environmental stimulus that is sensed by AdeS, as a number of conditions, including chlorhexidine shock, can also up-regulate *adeAB* expression in ATCC 17978 [54-57]. Instead, AdeS may respond to stimuli such as solutes that are excreted and accumulate in the periplasm when cells are subjected to various stressors including AdeAB substrates.

This study revealed unique interactions between pentamidine and a number of carbon sources which significantly alter the resistance profile to pentamidine for *A. baumannii* ATCC 17978 and derivatives. Biolog phenotypic arrays identified a number of succinic acid derivatives that also allowed respiration in the presence of a lethal concentration of pentamidine (Fig 4). Interestingly, when Δ*adeRS* utilized malonic acid as the sole carbon source, a potent inhibitor of the succinate dehydrogenase complex, cells were also able to respire at increased pentamidine concentrations (Fig 4). In Gram-negative bacteria, the mechanism of action for pentamidine is thought to primarily occur via binding to the lipid-A component of the outer membrane and not through inhibition of an intracellular target [23, 58]. Therefore, it seems unlikely that pentamidine interferes with enzymatic functions like that of succinate dehydrogenase, and instead the compounds which showed an increase in pentamidine resistance may contribute to either chelation of the compound or provide protective mechanisms that reduce binding to the lipid-A target. Interestingly, a 1956 study [59] identified that in the presence of α-ketoglutaric or glutamic acid, resistance to a lethal concentration of pentamidine could be achieved in *E. coli,* leading to the proposition that pentamidine interferes with the transaminase reaction in glutamic acid production. However, our study shows that glutamic acid does not affect pentamidine resistance (S4 Table), inferring that a different mechanism might be responsible for the increase in pentamidine resistance in *A. baumannii.*

Iron is an essential metal serving as a co-factor in numerous proteins involved in redox chemistry and electron transport. It is generally a limited resource for pathogenic bacteria such as *A. baumannii* where it is important for virulence and disease progression [6, 50]. Our studies showed that altering iron levels also had a marked effect on resistance to pentamidine. This is in agreement with the observation that citric and gluconic acids can act as iron-chelating agents [60, 61] and respiration activity in the presence of these compounds at the pentamidine concentrations tested were significantly altered (S3 Fig). Many clinically relevant antimicrobials have shown to be affected by the presence of iron, including compounds belonging to tetracycline, aminoglycoside and quinolone classes [22]. Activity of these compounds can be altered by numerous factors, including the formation of stable complexes which can affect drug efficacy or have unfavourable effects on patient health.

## Conclusions

Overall, this is the first study which has demonstrated that the AdeAB system in *A. baumannii* ATCC 17978 provides intrinsic resistance to a subset of dicationic compounds, and efflux of these compounds via AdeAB is directly regulated by the AdeRS TCSTS. RNA-seq identified that deletion of *adeRS* produced significant changes in the transcriptome where our results support the notion that strain specific variations are apparent [16]. We have provided evidence that in ATCC 17978, AdeRS is directly responsible for the activation of *adeAB* gene expression, as *ΔadeRS* failed to increase expression upon subjection to one of the pumps newly identified intrinsic substrates, pentamidine. It will be of interest to assess whether these dicationic compounds also extend as substrates towards AdeAB(C) pumps present in other *A. baumannii* isolates and if expression levels of *adeAB(C)* upon exposure to these substrates and other potential stressors also occur. This information may help to identify the stimulus that activates the AdeRS TCSTS. We have also demonstrated for the first time that for pentamidine to exert its antibacterial effect in *A. baumannii,* a dependence on the availability of iron is required, and that growth in the presence of selected carbon sources has a profound effect on its resistance.

## Acknowledgments

We would like to thank Professor Bryan Davies from the Department of Molecular Biosciences at the University of Texas for providing the plasmid pAT04.

## Supporting information

**S1 Fig. Validation of RNA-sequencing results.** The transcriptomic results obtained by RNA-sequencing were validated by qRT-PCR analysis. The level of nine genes that displayed differential expression or remained essentially unchanged between *ΔadeRS* and WT ATCC 17978 were chosen for comparison. Expression levels for qRT-PCR experiments were corrected to those obtained for *GAPDH* (ACX60_05065) prior to normalisation against WT ATCC 17978 transcriptional levels. Grey and black bars represent values obtained from RNA-seq and qRT-PCR results, respectively. Differential expression between *ΔadeRS* and WT ATCC 17978 are given in Log_2_-values.

**S2 Fig. Structures of dicationic antimicrobial compounds to which *ΔadeRS, ΔadeA, ΔadeB* and *ΔadeAB* deletion mutant derivatives showed a decrease in resistance when compared to WT ATCC 17978.** Compounds include (a) pentamidine, (b) DAPI and (c) chlorhexidine. For pentamidine and chlorhexidine, the cationic nitrogenous groups are separated by a long carbon chain, forming symmetrical compounds, whereas, DAPI lacks this long linker and is asymmetric

**S3 Fig. Comparative analysis of kinetic response curves obtained from Biolog PM plates for *ΔadeRS* cells untreated and subjected to increasing concentrations of pentamidine.** Respiration of ATCC 17978 *ΔadeRS* cells in the presence of pentamidine (8, 16, 32, or 64 mg/L) against untreated control cells are shown for (a) Biolog PM01 plates and (b) Biolog PM02A plates. Respiration activity for both plates were monitored in IF-0 (Biolog, Inc.) liquid medium for 72 h at 37°C. The curve in each well represents the colour intensity of a redox-active dye (*y* axis) over time (x axis: 72 h). Respiration of *ΔadeRS* cells are shown in red (control), green (under different concentrations of pentamidine), and yellow (depicts the regions of respiratory overlap). Black numbered squares represent carbon sources which decreased resistance to pentamidine (1, D-gluconic acid; 2, citric acid).

**S4 Fig. Pentamidine resistance is affected by the concentration of iron within the growth medium. Resistance to pentamidine was assessed by disc diffusion assays in Mueller-Hinton agar (MHA) for ATCC 17978 and ΔadeRS, ΔadeA, ΔadeB and ΔadeAB deletion derivatives.** Zones of clearing were compared to iron rich conditions from the addition of ferrous sulphate (FeSO_4_) at the final concentrations of 2.5 and 5 mM and iron-chelated conditions obtained by the addition of 2’,2’ dipyridyl (DIP) at the final concentrations of 100 and 200 μM in MHA. Images displayed are a representative of the typical results obtained.

**S1 Table. Strains and plasmids used in the study.**

**S2 Table. Primers used in this study.**

**S3 Table. Genes significantly up- and down-regulated (≥ 2-fold) in *AadeRS* compared against the parent ATCC 17978 (CP012004) by RNA-seq methodologies.**

**S4 Table. Zones of clearing obtained from growth on M9 minimal medium with the addition of different carbon sources after exposure to pentamidine.**

